# PRESENCE OF A LARGE DASYPODIDAE GRAY, 1821 FROM THE CUVIERI CAVE, EASTERN BRAZIL

**DOI:** 10.1101/2025.01.08.631973

**Authors:** Artur Chahud

## Abstract

Cingulata represent a group of Xenarthra characterized by the presence of a carapace composed of bony plates (osteoderms). The Cuvieri Cave, located in the Lagoa Santa karst complex, presents a relevant quantity of vertebrate fossils, including extinct and extant species. Although several paleontological studies have been carried out, the material was reviewed from a taxonomic point of view, including Xenarthra material. Two osteoderms, initially attributed to the genus *Euphractus*, were found in Pleistocene deposits, but it was observed that the specimens did not belong to this genus. The aim of this study is to present the revision of these two specimens, now recognized as the first evidence of a large member of the family Dasypodidae in the Cuvieri Cave. The specimens were found in levels where both extant and extinct species from the Lagoa Santa region were found, but due to the paleoenvironmental changes of the Pleistocene-Holocene transition in Lagoa, it was not possible to determine the environment in which the specimen lived.

## INTRODUCTION

The Superorder Xenarthra comprises a clade of mammals of South American origin that includes two orders: Cingulata (armadillos) and Pilosa (anteaters and sloths). The earliest fossil records of this Superorder date back to the Paleocene (Fernicola et al., 2021), with its distribution restricted to the South American continent until the Neogene, a period during which the Great American Biotic Interchange (GABI) occurred, enabling these species to disperse into North America (Shapiro et al., 2015).

Cingulata represents a group of Xenarthra characterized by the presence of a carapace composed of bony plates (osteoderms), covering the back, flanks, the upper region of the head, and the tail. In current armadillo species, the dorsal carapace is segmented into transverse bands; in the scapular and pelvic regions, these bands are fixed, while the intermediate bands between these areas are movable. This morphological arrangement allows individuals of this clade to curl up into a spherical shape to some extent, as exemplified by the genus *Tolypeutes* Illiger, 1811 (three-banded armadillo) (Paula Couto, 1979).

Fossils of Cingulata are frequently found in Quaternary deposits in South America, with records of the genera *Zaedyus* Ameghino, 1889, *Chaetophractus* Fitzinger, 1871, *Chlamyphorus* Harlan, 1825, *Calyptophractus* Fitzinger, 1871, *Priodontes* F. Cuvier, 1825, *Tolypeutes* Illiger, 1811, *Cabassous* McMurtrie, 1831, *Euphractus* Wagler, 1830, and *Dasypus* Linnaeus, 1758 (Delsuc et al., 2016). The latter three were identified in paleontological and archaeological sites dated to the Holocene from southeastern Brazil (Ameghino, 1907; Paula-Couto, 1973; 1975; 1979; Chahud, 2021; Chahud et al. 2021; 2022).

The Cuvieri Cave, located in the karst complex of Lagoa Santa, has been the subject of multiple studies involving dating, speleology and paleontology (Alvarenga et al., 2008; Hubbe et al., 2011; Mayer et al., 2016; Haddad-Martim et al., 2017; Chahud, 2020a, 2020b, 2020c; Chahud et al., 2020; Chahud & Okumura, 2021a; 2021b).

Recently, Chahud (2021) identified and described a specimen of *Euphractus* Wagler, 1830, from the Holocene of Cuvieri Cave. The author also reported that two osteoderms, initially attributed to this genus, were found in Pleistocene deposits, being considered evidence of the presence of *Euphractus* during that period. However, after analysis and revision, it was concluded that the specimens in question do not belong to the genus *Euphractus*. The objective of this study is to present the revision of these two specimens, now recognized as the first evidence of a large member of the family Dasypodidae in Cuvieri Cave.

## MATERIAL AND METHODS

The Cuvieri Cave is located at UTM coordinates: 23K 603756E and 7846105S, in the state of Minas Gerais, southeastern Brazil (Figure 1). This cave features semi-consolidated deposits, which facilitated the application of detailed excavation techniques and allowed the acquisition of reliable stratigraphic data.

**Figure 1.**
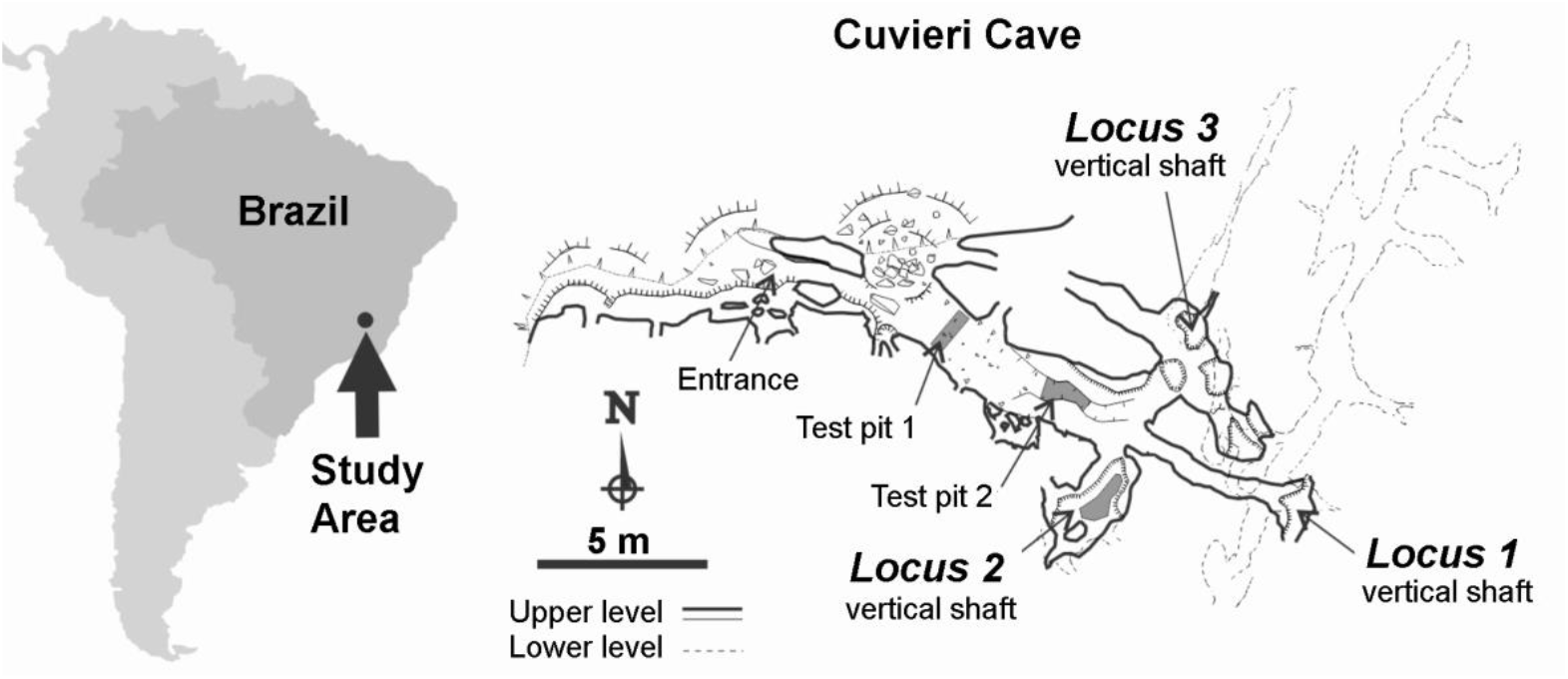
Geographic location of the study area and of Cuvieri Cave showing the position of *Loci* 1, 2, and 3 (map courtesy of Alex Hubbe and Grupo Bambuí for Speleological Research).

The cave has two passages: a larger, obstructed one and a smaller, open one, measuring 1.5 m in height and 1 m in width, through which the interior of the cave can be accessed. Inside, there are three small vertical cavities, forming natural traps called *Locus* 1, *Locus* 2, and *Locus* 3 (Figure 1), with depths of 16 m, 4 m, and 8 m, respectively.

The osteoderms presented in this study come from *Locus* 3 and are close both stratigraphically and in location within the cave. No other associated bone parts were found, suggesting that these specimens belonged to an individual that was not fully preserved or that they were transported into *Locus* 3 from an external area. The specimens show few signs of abrasion, with one of them exhibiting fragmentation.

The age of the deposits from *Locus* 3 was considered late Pleistocene, based on radiocarbon dating of specimens found near the surface layers, with minimal reworking and transportation, as well as speleothem samples (Haddad-Martim et al., 2017).

For the identification of the materials, comparisons were made with specimens from scientific collections such as the “Renato Kipnis Collection” from LEEH (Laboratory of Human Evolutionary Studies, Department of Genetics and Evolutionary Biology, Institute of Biosciences, University of São Paulo), the mammal collection of the Museum of Zoology of the University of São Paulo, and consultations with works by Paula Couto (1979), Pitana & Ribeiro (2007), Oliveira & Pereira (2009), Castro et al. (2013) Delsuc et al. (2016), Feijó & Cordeiro Estrela (2016), Feijó et al. (2018),.

The specimens were collected in 2005, as part of the thematic project “Origens e Microevolução do Homem na América: uma abordagem paleoantropológica”, coordinated by Prof. Dr. Walter Neves. They were later curated, numbered, and registered in the Laboratory of Human Evolutionary Studies.

## SYSTEMATIC PALEONTOLOGY

Superorder Xenarthra Cope, 1889

Order Cingulata Illiger, 1811

Superfamily Dasypodoidea Gray, 1821

Family Dasypodidae Gray, 1821

Figure 2

**Figure 2.**
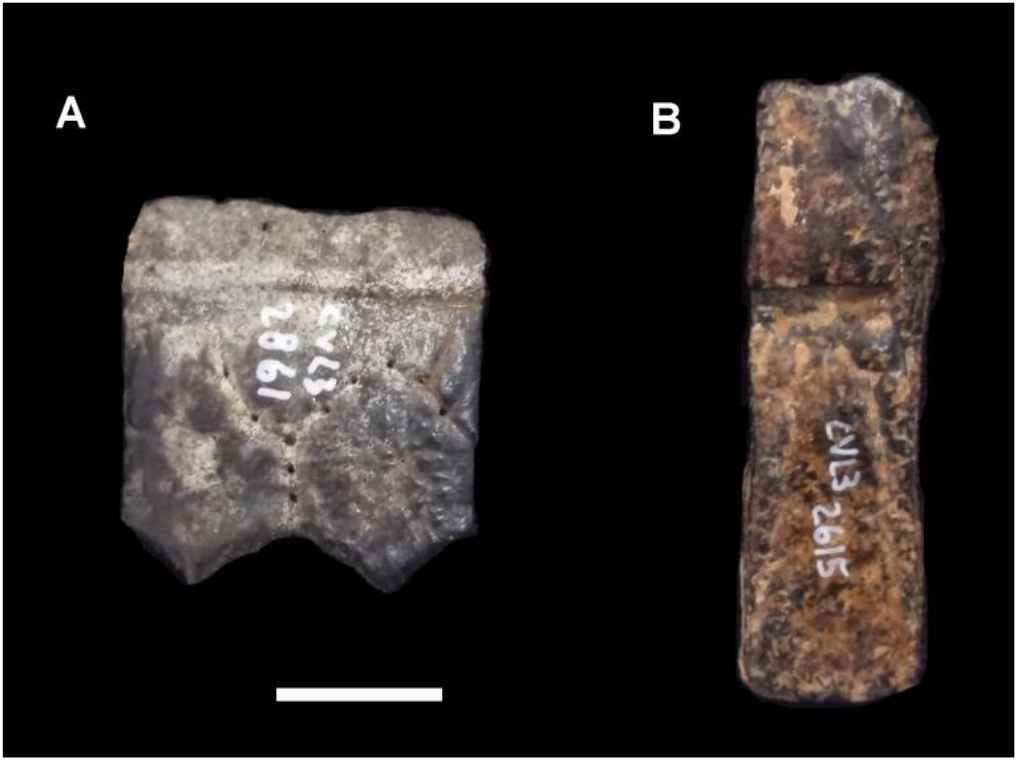
Osteoderms of Dasypodidae from *Locus* 3 of Cuvieri Cave. A) CVL3-2861 and B) CVL3-2615. Scale 10 mm.

### Material

The analyzed specimens are represented by two disarticulated osteoderms, identified as CVL3-2615 and CVL3-2861.

### Remarks

The osteoderms, although originating from different positions on the carapace, likely belong to the same individual, as no other specimens were found, and they are located only a few centimeters apart.

The CVL3-2861 specimen corresponds to the front edge portion of the carapace. It has a quadrangular shape, with two hexagonal contour structures separated by grooves and small foramina. The edge is marked by a slightly concave, whitish band. Its dimensions are 23 mm wide and 23.5 mm long, with a thickness of approximately 5 mm.

The CVL3-2615 specimen belongs to the mobile belt region of the carapace, with an elongated rectangular shape and, on the surface, it shows foramina and two prominent, diverging lateral grooves, forming a “V” shape. This osteoderm, divided into two parts, has a smooth, thicker articulation area in the anterior part and diverging grooves in the posterior part. It is considerably thinner than CVL3-2861, with a thickness of less than 1 mm, and its dimensions are 37 mm long and 12 mm wide.

### Discussion

Initially, due to the size of the specimens and evidence of surface abrasion, they were classified as belonging to the living armadillo *Euphractus*, both by the collectors and by Chahud (2021). However, the absence of central rugosities and the “V” shape of the diverging grooves in specimen CVL3-2615 are features observed in members of the family Dasypodidae.

Although they have larger dimensions than current Dasypodidae, the specimens display superficial ornamentation and smaller dimensions compared to large Cingulata specimens, such as the Pampatheriidae, which have similar osteoderms (Pereira Ferreira et al., 2025) or Glyptodontinae.

The observed structures and thickness of specimen CVL3-2861 are not found in living Dasypodidae specimens in the Lagoa Santa region, but they were found in extinct specimens of *Propraopus* (Pitana & Ribeiro, 2007) and the currently extinct species *Dasypus punctatus* (Castro et al., 2013). The mobile osteoderm CVL3-2615 has a size compatible with those observed in *Propraopus* and *D. punctatus*.

Despite the similarity and proportion with the genus *Propraopus*, specimen CVL3-2615 also has an external morphology similar to those found in the genus *Dasypus*, both current and extinct, possibly representing the species *D. punctatus*, observed in the southeastern region of Brazil (Castro et al., 2013). Specimen CVL3-2861 showed characteristics such as a 5 mm thickness and the edge, which were not observed in living specimens of *Dasypus* in the Lagoa Santa region.

In neither specimen was it possible to determine the number of foramina to define if they belonged to the species *Dasypus punctatus*, and therefore the association with this species cannot be characterized. However, they also did not present diagnostic characteristics of the genus *Propraopus*, and therefore it was decided not to classify them at the genus or species level.

## ASSOCIATED FAUNA

The two specimens were found in layers containing remains of animals typical of the current fauna, such as the cervids *Mazama americana* and *Subulo cf. gouazoubira* (Chahud et al., 2023a), the large feline *Panthera onca* (Chahud & Okumura, 2021a), the leporid *Sylvilagus sp*. (Chahud et al., 2020; Chahud & Okumura, 2022), and the large rodents *Cuniculus paca* and *Dasyprocta sp*. (Chahud, 2020a; 2022). There were also remains of extinct or indeterminate species of current families, such as Cuniculidae, Tapiridae, and Tayassuidae (Mayer et al., 2016; Chahud et al., 2023a; 2023b). However, in Cuvieri Cave, extinct species of the Pleistocene megafauna, such as ground sloths and the saber-toothed tiger *Smilodon populator*, were also observed in more recent layers (Hubbe et al., 2011; Chahud, 2020b).

Although the living species in the region are present in the deposit, it is not possible to consider that the environment was the same as the current one, as previous studies have observed that during the late Pleistocene in Lagoa Santa, some significant paleoenvironmental changes occurred, especially near the transition between the Holocene and Pleistocene periods (Ledru, 1993; Ledru et al., 1996).

## CONCLUSIONS

Contrary to what was identified in previous studies, the Pleistocene specimens did not belong to an *Euphractus* specimen but to a large Dasypodidae individual, possibly attributed to the genus *Propraopus* or the species *Dasypus punctatus*. Both were found in Lagoa Santa and various locations in Brazil.

The specimens were found in deposits dated to the late Pleistocene and associated with specimens of the current fauna of the Lagoa Santa region, as well as in older deposits than those in which extinct representatives of the regional megafauna were identified. However, due to significant environmental changes during this period, it is not possible to precisely define the paleoenvironment in which the specimen lived.

## ACKNOWLEDGEMENTS

The author thanks Dr. Maria Mercedes Martinez Okumura, responsible for LEEH (Laboratory for Human Evolutionary Studies), Department of Genetics and Evolutionary Biology, Institute of Biosciences at the University of São Paulo, for allowing the preparation of fossils in the laboratory.

